# Unsupervised nonmotor learning in the human cerebellum

**DOI:** 10.64898/2026.07.22.740142

**Authors:** Juliana E. Trach, Yiran Ou, Nicholas B. Turk-Browne, Samuel D. McDougle

## Abstract

A core motor function of the cerebellum is error-based learning: it uses prediction errors—mismatches between predicted and actual sensory feedback—to refine actions. One provocative hypothesis is that the cerebellum performs similar computations in cognitive domains. Here, we ask whether error-based learning computations in the cerebellum extend to a passive statistical learning task that requires neither action nor decision-making. Human participants viewed sequences of visual stimuli with varying probabilistic stimulus-stimulus transitions while undergoing fMRI. Subjects reported no explicit knowledge of the transitional probabilities after the scan. Nonetheless, putative cognitive regions of the cerebellum encoded prediction errors during statistical learning. Moreover, error signals in the cerebellum were encoded in a different manner than error signals in the anterior hippocampus, a well-known substrate of statistical learning: cerebellar activity covaried with trial-by-trial prediction errors that evolved slowly over time (echoing cerebellar computations in motor learning), whereas the hippocampus responded in a more fixed manner to the underlying transition structure. Computational modeling formalized this dissociation, with cerebellar activity explained by a delta-rule model that incrementally adjusted predictions based on recent experience, and hippocampal activity characterized as a form of Bayesian updating of a transition distribution held in memory. Our findings demonstrate that the cerebellum encodes prediction errors during passive nonmotor learning, and thus that it likely plays a domain-general role in error-based learning.

**Significance:** The cerebellum is known to support motor learning by using prediction errors to refine behavior. Here, we test whether these computations extend to nonmotor learning contexts that require neither action nor decision-making. We report the discovery of prediction error signals in the cerebellum during statistical learning, an unsupervised process that extracts regularities from the sensory environment. These findings support a domain-general role for the cerebellum in error-based learning, offering a possible explanation for why cerebellar damage is implicated in such wide-ranging outcomes, from impaired motor coordination to complex disorders like autism.

## Introduction

Supervised motor learning is one of the cerebellum’s most widely studied functions (1–5). In this domain, the cerebellum generates predictions about the sensory consequences of actions and uses discrepancies between predicted and actual feedback (i.e., prediction errors) to refine behavior (5–8). The cerebellum is an ideal substrate for this type of error-based learning because it contains two parallel input pathways. One pathway (mossy fiber inputs) carries high-dimensional state information (e.g., motor commands and sensory feedback), while the other pathway (climbing fiber inputs) carries error information (i.e., the discrepancy between the expected outcome and the current state) (9). The segregation of these signals allows the error information to drive learning in Purkinje cells without corrupting the state information in mossy fibers. This stereotyped circuit motif has led to proposals that the cerebellum may perform similar computations across motor and nonmotor domains (10, 11). Inspired by this idea and growing interest in “cerebellar cognition” (11–20), we ask whether the established role of the cerebellum as a prediction-error learning system generalizes to a form of nonmotor learning: visual statistical learning (SL) (21). Establishing the existence of cerebellar signals related to this kind of passive, nonmotor learning would support the idea that the cerebellum encodes generalized prediction errors across domains.

Recent work has begun to explore nonmotor predictive learning computations in the cerebellum during reinforcement learning (RL) (22–27). RL describes how behavior is modified over time through feedback about the rewarding or aversive consequences of actions (28, 29). During RL, an agent makes predictions about how rewarding particular actions will be, and uses discrepancies between predicted and actual reward (i.e., reward prediction errors) to update internally represented action values. Neural correlates of RL have been identified across the cerebellar circuit in model organisms and humans (24, 25, 27, 30) and subtle RL-related behavioral deficits have been observed in humans with cerebellar degeneration (31, 32). Human neuroimaging research has highlighted functional areas spanning the border of lobules Crus I and II (26) as the primary loci of cerebellar RL signals, consistent with the cognitive “S2” region in state-of-the-art functional maps of the human cerebellum (33).

One complication with studying RL in the cerebellum is that such tasks typically require or induce motor responses (e.g., button presses in human research, lever pulls or licking in model organisms). Thus, the encoding of RL signals in the cerebellum might be dependent on the action-based nature of RL, qualifying claims about “domain-general” cerebellar learning functions beyond motor behavior. Here we avoid these issues, asking whether prediction error signals are present in the cerebellum during a passive, visual SL task that does not require action or even decision-making.

SL involves extracting regularities from sensory inputs over space and time (34–36). SL is a ubiquitous “unsupervised” learning process that is functional from early infancy (35, 37), allowing organisms to learn about the structure of the environment without taking actions or evaluating feedback. For example, you might expect a knock at your door when you hear your dog bark because your dog always barks when someone arrives in the driveway. There is a robust neuroscience literature on SL that focuses on medial temporal lobe (MTL) structures, like the hippocampus (Hpc) and parahippocampal cortex. This work has demonstrated that the Hpc is necessary for effective SL (38, 39). More specifically, SL is thought to leverage the monosynaptic pathway that connects entorhinal cortex to hippocampal subfield CA1 (40, 41), localizing SL signals in Hpc to its anterior portion. In terms of prediction-error activity during SL, CA1 has been shown to exhibit elevated responses when a statistical expectation is violated (42), along with other MTL regions like parahippocampal cortex (42), and perirhinal cortex (43, 44). To our knowledge, cerebellar contributions to SL remain unexplored.

In the present study, we use model-based whole-brain fMRI to test whether and how the human cerebellum encodes prediction errors when sensory expectations are violated during visual SL. Further, we ask how cerebellar prediction errors during SL are modulated by the temporal interval between sensory events, echoing constraints observed in motor learning (45–49) and RL (26). Finally, we take a computational approach to formalize different potential mechanisms underlying SL, allowing us to contrast cerebellar versus hippocampal prediction-error signals. Overall, our study provides a strong test of hypothesized domain-general learning functions of the cerebellum by examining prediction-error signals during purely unsupervised visual SL.

## Results

### Unsupervised visual statistical learning

Twenty participants completed a visual SL task while undergoing fMRI. Each run included four unique, colorful fractals used in prior SL studies, which were presented one at a time (**Figure 1a**) (50). We manipulated the transitional probability between fractals such that each fractal was typically followed by one other fractal (common transition, probability 0.75), rarely followed by a different fractal (rare transition, probability 0.25), and never followed by itself nor the other fractal in the set (**Figure 1b**).

**Figure 1.**
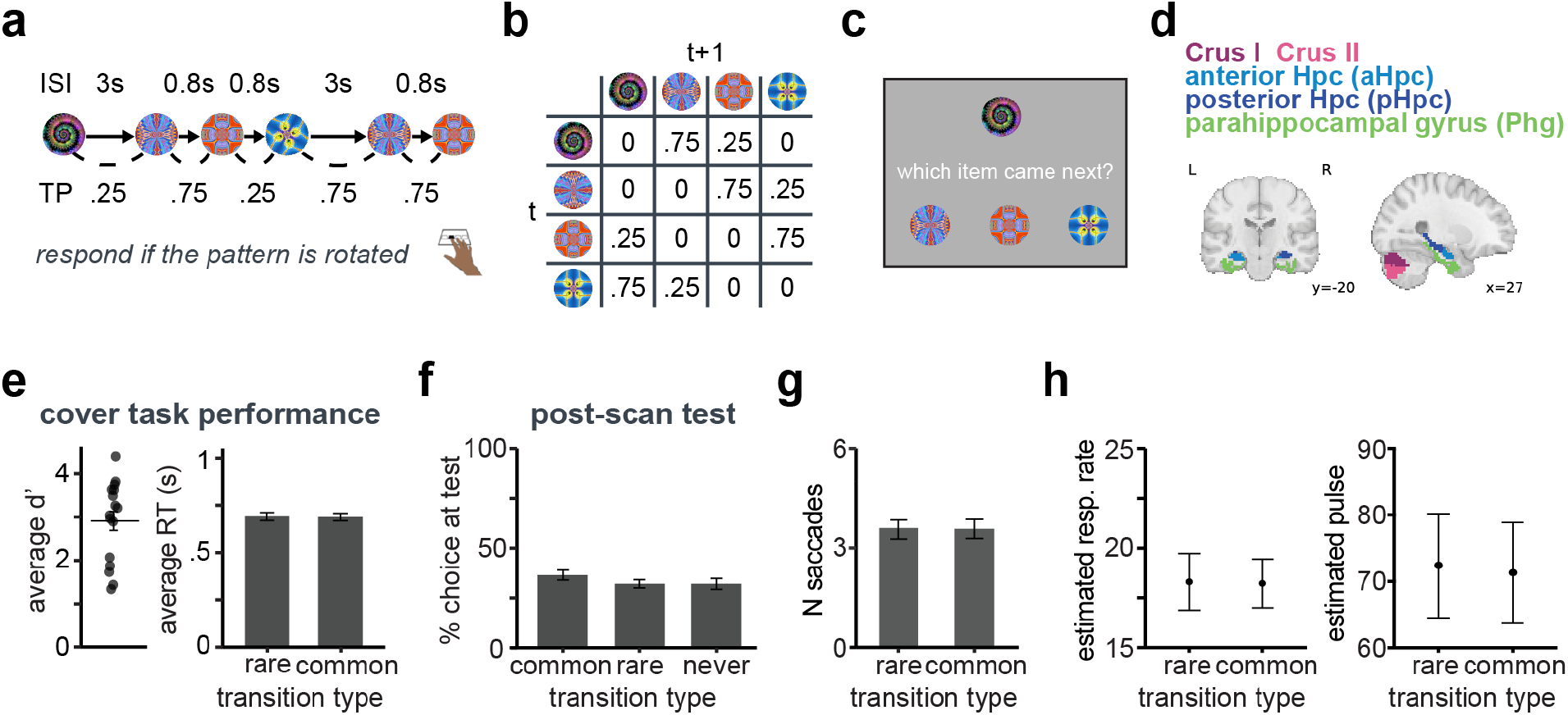
Task and behavior. a) Task schematic. Subjects passively viewed a sequence of fractal images. The cover task was to detect when a fractal was rotated from its upright orientation (e.g., the fourth fractal in this example) and press a button. b) Example transition matrix for a block’s fractal sequence (i.e., column = fractal on trial t+1, row = fractal on trial t). Note the presence of common (0.75) and rare (0.25) transition types. c) Post-scan explicit test example trial. d) Anatomical ROIs for the main analyses. e) Average d-prime and average RT on cover task. f) Performance on post-scan test. g) Gaze behavior (number of saccades) by transition type. h) Respiration and pulse by transition type.

In supervised motor learning and RL, cerebellar learning is temporally sensitive and blunted by suprasecond delays between key events, such as a predictive sensory cue and an eyepuff in Pavlovian eyeblink conditioning (48). We tested whether putative cerebellar prediction errors are time-sensitive in SL by manipulating the interstimulus interval (ISI) between fractals; short-delay trials were preceded by 0.8-s ISIs and long-delay trials were preceded by 3-s ISIs. Finally, participants were instructed to press a button if the presented fractal was rotated from its original orientation as a cover task unrelated to the transition structure (**Figure 1a**).

Participants performed the cover task well (**Figure 1e, left panel**; average d-prime: *M* = 2.91, *SE* = 0.22) and had comparable RTs across rare versus common transitions (RT: *t*(16) = 0.26, 95% CI = [-0.02, 0.02], *p* = .795; **Figure 1e, right panel**). After scanning, we tested participants for explicit knowledge of the transitional structure (**Figure 1c**). At a group level, participants did not show evidence of explicit knowledge of the structure **(Figure 1f**; common > rare: *t*(19) = 1.17, 95% CI = [-0.03, 0.12], *p* = .257; common > never: *t*(19) = 0.92, 95% CI = [-0.06, 0.14], *p* = .371), consistent with this form of SL being implicit (21, 51, 52).

Eye movements (**Figure 1g**) and physiological signals (**Figure 1h**) can influence cerebellar activity. However, we observed no relationships between these measures and key stimulus events, such as rare versus common transitions (N saccades: *t*(17) = -0.02, 95% CI = [-0.08, 0.08], *p* = .984; respiration rate: *t*(12) =0.39, 95% CI = [-0.37 0.53], *p* = .704; pulse: *t*(10) = 1.78, 95% CI = [-0.27, 2.43], *p* = .106).

### Cognitive regions of cerebellum track incremental prediction error, rather than transition type

We considered two types of prediction errors the brain may compute during SL. First, we examined how the brain responded to rare versus common transitions. Increased activity on a rare transition (relative to a common transition) is indicative of prediction-error processing (“mismatch” effects) (53, 54). We contrasted activity on rare versus common transitions using a general linear model (GLM).

Second, we adopted a simple delta-rule learning model to examine incremental prediction errors (28) (**Figure 2c**). This type of model, which gradually updates expectations based on the discrepancy between predicted and expected sensory inputs, approximates computations the cerebellum performs in supervised motor learning (1, 5, 6, 8, 9). We used a fixed learning rate of 0.005 as we did not have explicit learning behavior to fit. Then we simulated a timeseries of trial-wise prediction errors for each participant and entered these as parametric regressors in an fMRI GLM analysis.

**Figure 2.**
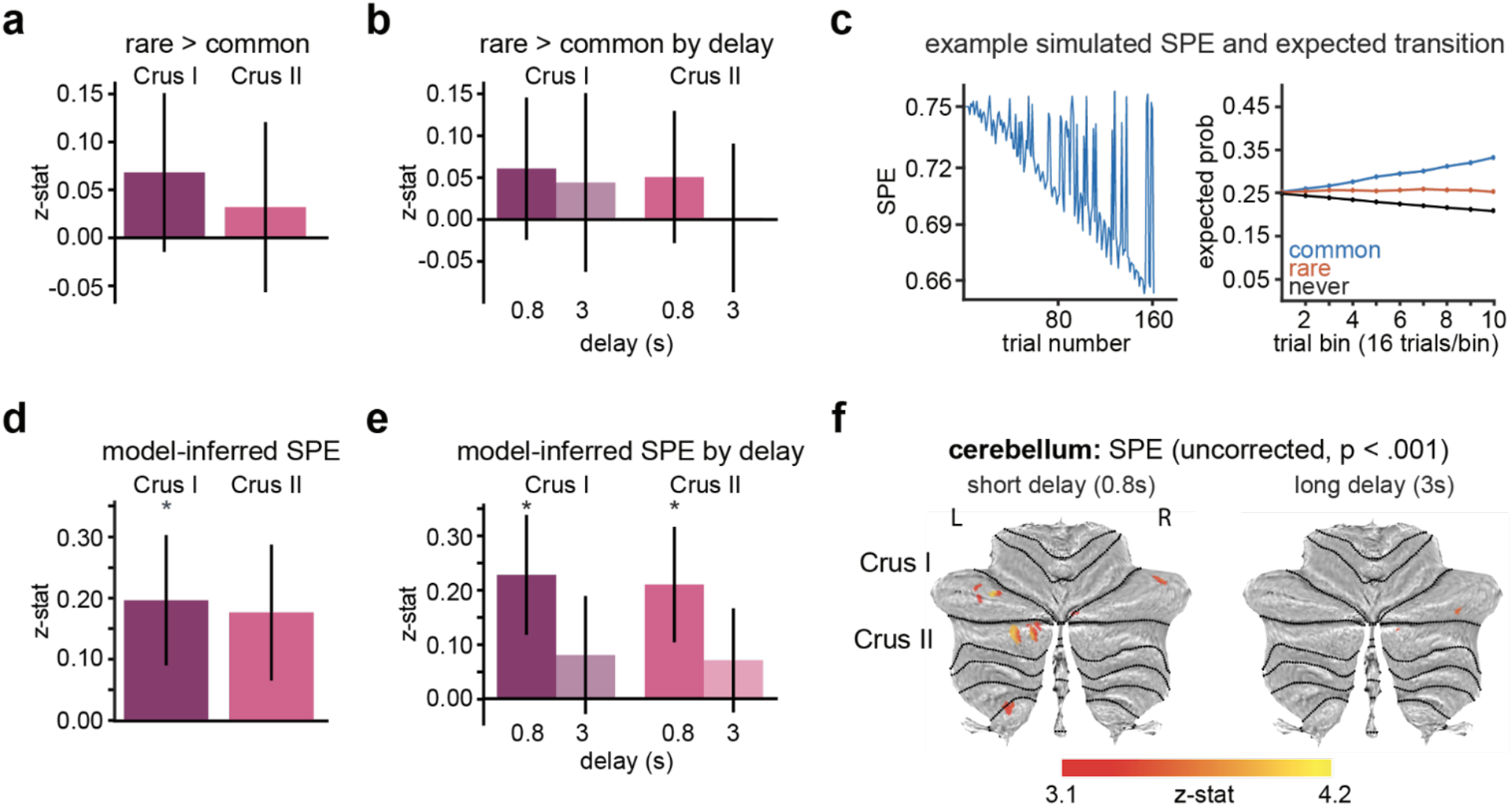
Cerebellar results. a) Contrast of activity on rare > common transitions in cerebellar lobular ROIs. b) Same as (a), by delay condition. c) Example timeseries of model-predicted sensory prediction errors (SPEs) and incremental learning of the probability of each transition bigram. d) SPE response in cerebellar ROIs. e) SPE response in cerebellar ROIs by delay condition. f) SPE response in whole cerebellum (uncorrected, p < .001). * p < .05. Error bars = S.E.M.

We focused analysis on Crus I and II lobules in the cerebellum (**Figure 1d**), which are generally associated with cognitive functions in the cerebellum (26, 33), and on the hippocampus given its known involvement in SL.

We did not find evidence that the cerebellum tracked transition type across all trials (rare > common: Crus I: *M* = 0.07 95% CI = [-0.08, 0.23], *p* = .348; Crus II: *M* = 0.03, 95% CI = [-0.14, 0.19], *p* = .754; **Figure 2a**), at short delays (Crus I: *M* = 0.09, 95% CI = [-0.08, 0.25], *p* = .304; Crus II: *M* = 0.05, 95% CI = [-0.09, 0.19], *p* =.496), or at long delays (Crus I: *M* = 0.01, 95% CI = [-0.17, 0.20], *p* = .934; Crus II: *M* = -0.002, 95% CI = [-0.17, 0.16], *p* = .984; **Figure 2b**).

In contrast, we did observe significant incremental prediction error encoding in Crus I (**Figure 2d**; *M* = 0.196, 95% CI = [0.02, 0.40], *p* = .032), and marginal effects in Crus II (*M* = 0.18, 95% CI = [-0.02, 0.40], *p* = .076). These effects were driven by robust prediction error coding across short delays (**Figure 2e**) in both Crus I (*M* = 0.23, 95% CI = [0.03, 0.44], *p* = .022) and Crus II (*M* = 0.21, 95% CI = [0.03, 0.41], *p* = .018), and these effects were not robust at long delays in Crus I (*M* = 0.08, 95% CI = [-0.13, 0.28], *p* = .484) or Crus II (*M* = 0.07, 95% CI = [-0.12, 0.26], *p* = .464). Subsequent exploratory analysis of the whole cerebellar volume revealed short-delay prediction error encoding in clusters spanning Crus I and II, as well as a smaller cluster around the boundary between lobule IX and VIIIb (**Figure 2f**).

### Hippocampus responds to surprising transitions rather than incremental prediction errors

The MTL showed a different pattern of results than the cerebellum. We used ASHS to obtain participant-specific anatomical segmentations of ROIs: anterior hippocampus (aHpc), posterior hippocampus (pHpc), parahippocampal gyrus (Phg) (55). We found that aHpc responded to transition type, with higher activity to rare versus common transitions (*M* = 0.28, 95% CI = [0.07, 0.48], *p* = .004) (**Figure 3a**); pHpc (*M* = 0.11, 95% CI = [-0.05, 0.29], *p* = .200) and Phg (*M* = 0.11, 95% CI = [-0.003, 0.22], *p* = .060) were in the same numerical direction but not significant. This effect emerged with learning: we found the aHpc effect was robust in the second half of exposure (*M* = 0.37, 95% CI = [0.13, 0.61], *p* = .004) but not the first half (*M* = 0.08, 95% CI = [-0.08, 0.23], *p* = .300) (**Figure 3b**). Phg likewise showed a reliable effect in the second half (M = 0.12, 95% CI = [0.008, 0.25], p = .032) but not first half (M = 0.002, 95% CI = [-0.12, 0.13], p = .952); pHpc showed no reliable effects in second half (M = 0.12, 95% CI = [-0.07, 0.33], p = .252) or first half (M = 0.04, 95% CI = [-0.15, 0.22], p = .646).

**Figure 3.**
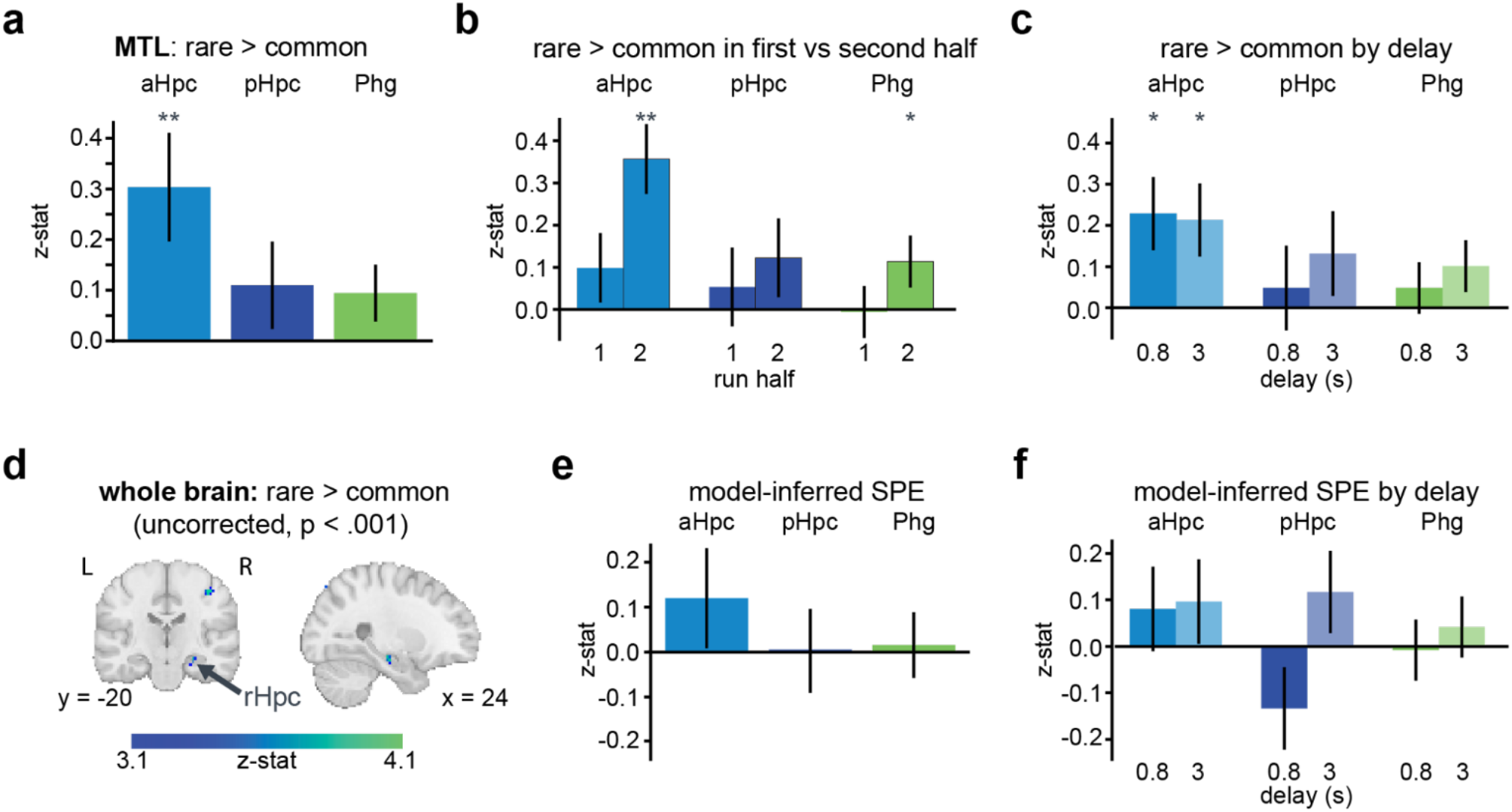
Medial temporal lobe results. a) Contrast of activity on rare > common transitions in MTL ROIs. b) Rare > common effect in MTL ROIs separately for first half of trials and second half of trials. c) Same as (a), by delay condition. d) Exploratory whole-brain results for rare > common transition contrast (uncorrected, p < .001). e) SPE response in cerebellar ROIs. f) SPE response in cerebellar ROIs by delay condition. ** p < .01; * p < .05. Error bars = S.E.M.

Prediction error responses in aHPC were consistent across short (*M* = 0.20, 95% CI = [0.01, 0.39], *p* = .034) and long delays (*M* = 0.19, 95% CI = [0.007, 0.37], *p* = .042). There was also a marginal effect at long delays in pHPC (*M* = 0.13, 95% CI = [-0.02, 0.27], *p* = .078), but not at short delays (*M* = 0.034, 95% CI = [-0.17, 0.22], *p* = .776) nor in Phg (short delays: *M* = 0.07, 95% CI = [-0.06, 0.20], *p* = .280; long delay: *M* = 0.08, 95% CI = [-0.02, 0.19], *p* = .132). Significant clusters in Hpc and other MTL structures were also evident in exploratory whole-brain analyses (**Figure 3d; Supplemental Figure 1**).

In contrast to our findings in the cerebellum, we did not find significant encoding of incremental model-based prediction errors in the MTL ROIs (**Figure 3e**; aHpc: *M* = 0.12, 95% CI = [-0.09, 0.35], *p* = .258; pHpc: *M* = 0.003, 95% CI = [-0.16, 0.19], *p* = .986; Phg: *M* = 0.02, 95% CI = [-0.13, 0.15], *p* = .838), even when separating short delays (**Figure 3f**; aHpc: *M* = 0.08, 95% CI = [-0.09, 0.27], *p* = .400; pHpc: *M* = - 0.13, 95% CI = [-0.29, 0.03], *p* = .126; Phg: *M* = -0.008, 95% CI = [-0.14, 0.12], *p* = .904) and long delays (aHpc: *M* = 0.10, 95h% CI = [-0.09, 0.29], *p* = .336; pHpc: *M* = 0.12, 95% CI = [-0.06, 0.30], *p* = .204; Phg: *M* = 0.04, 95% CI = [-0.07, 0.16], *p* = .506). Thus, aHpc exhibits prediction errors related to transition type, rather than delta-rule-based prediction errors seen in the cerebellum.

### Distinct computational algorithms for SL prediction errors in cerebellum versus hippocampus

The contrasting prediction error profiles in the cerebellum and hippocampus indicate a double dissociation, where cognitive regions of the cerebellum selectively encoded incremental, trial-by-trial SL prediction errors captured by a delta-rule model, whereas the hippocampus selectively tracked transition type. We further explored this dissociation by comparing the fit of a delta-rule learning model versus a Bayesian learning model to activity in our ROIs. In contrast to the delta-rule model, which generates incremental prediction errors, a Bayesian model accrues individual episodes over time and gains confidence in the underlying statistical structure of the environment (56, 57). In this type of model, a rare transition late in the task will only add confidence to the underlying learned transition model, while a delta-rule model will still insist that a strong update is needed.

We extracted BOLD timecourses from ROIs and fit a delta-rule and a Bayesian SL model to these timecourses to derive the best-fitting model parameters for each participant, and computed betas for each model using a cross-validation procedure. Hippocampal activity was better fit by the Bayesian model (**Figure 4a**; *z* = 2.58, *p* = .01) than the delta-rule model (*z* = 0.49, *p* = .627; delta-rule vs. Bayesian: *z* = 2.87, *p* = .004). In the cerebellum, the cross-validated betas numerically favored the delta-rule model over the Bayesian model (Crus I: delta-rule model: *z* = 2.24, *p* = .025; Bayesian model: *z* = 1.49, *p* = .135; Crus II: delta-rule model: *z* = 2.02, *p* = .044; Bayesian model: *z* = 1.08, *p* = .279), although the direct contrast between models was not reliable (Crus I: *z* = -0.86, *p* = .391; Crus II: *z* = -1.16, *p* = .247). Critically, the difference of differences in cross-validated betas between brain regions was significant (**Figure 4b**; Crus I vs. Hpc: *z* = -2.24, *p* = .025; Crus II vs. Hpc: *z* = -2.54, *p* = .011), thus providing support for the hypothesis that the cerebellum and hippocampus employ qualitatively different learning algorithms.

**Figure 4.**
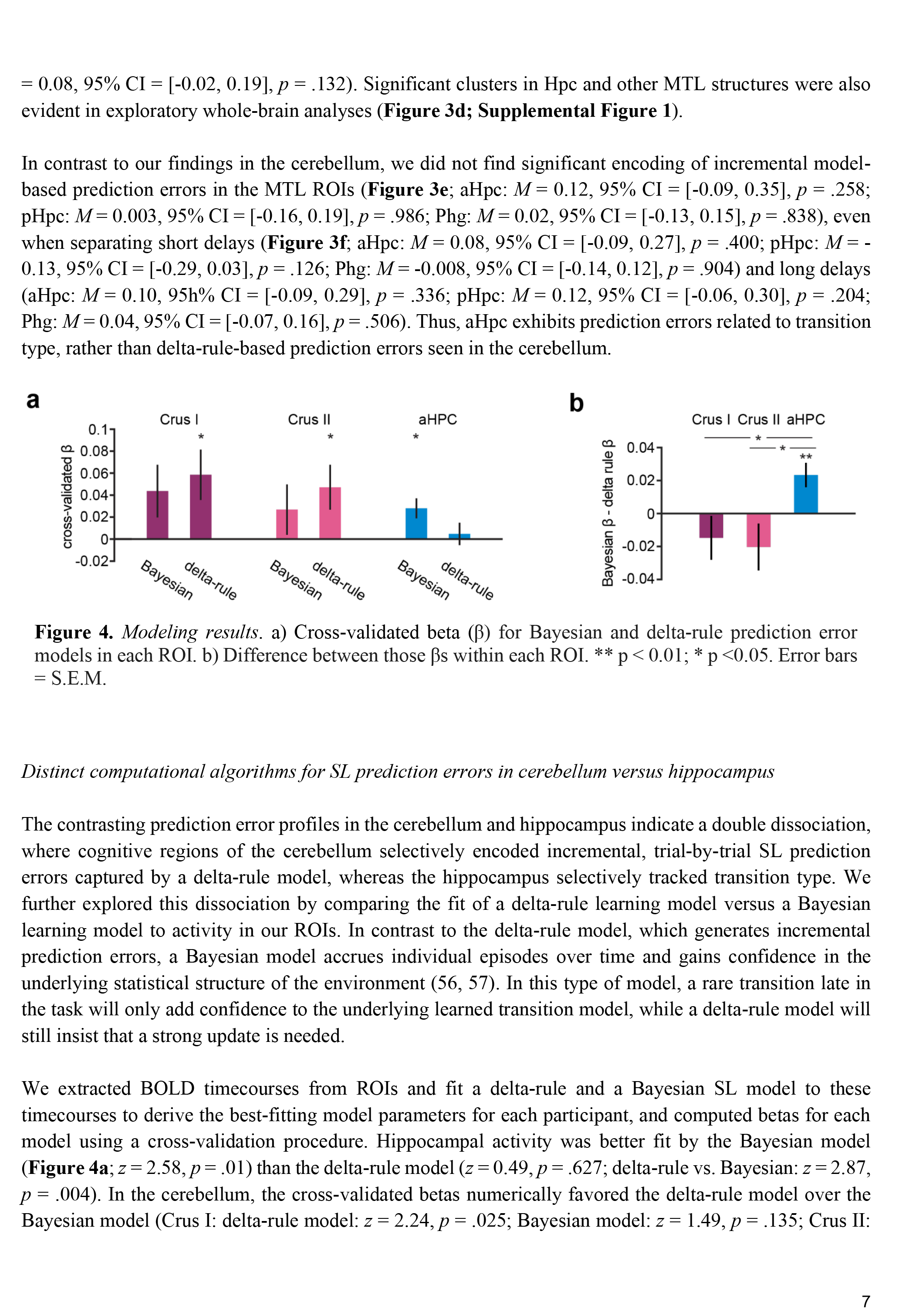
Modeling results. a) Cross-validated beta (β) for Bayesian and delta-rule prediction error models in each ROI. b) Difference between those βs within each ROI. ** p < 0.01; * p <0.05. Error bars = S.E.M.

## Discussion

We asked whether the cerebellum’s role in error signaling may generalize beyond supervised motor learning. In support of this possibility, we observed prediction error signals in the human cerebellum during unsupervised SL. These results expand upon recent work documenting RL signals in the cerebellum (23, 26, 27), to a novel context that required no action, decision-making, nor even feedback processing. Our findings suggest that cerebellar learning may not always require action, and that error-based learning may be a domain-general function of the cerebellum.

Cerebellar activity tracked sensory prediction errors derived from a standard delta-rule model that incrementally updates internal predictions trial-by-trial. Notably, foundational models of cerebellar motor learning (6–8) perform an analogous form of gradual learning, as do classic models of classical conditioning used to explain behavior in cerebellar-dependent conditioning tasks (58). These results suggest that the cerebellum may deploy a shared computational motif across supervised motor learning (1), RL (26), and unsupervised SL. Furthermore, the circuit-level dynamics of this process are well-characterized in motor learning (1) and have recently been argued to extend to RL (24), where error signals carried in climbing fibers drive plasticity in Purkinje cells and gradually alter internal predictions. Future work with invasive neurophysiology will be essential to understand whether SL may operate in a similar manner at the circuit level.

Our results also suggest that the cerebellum and the hippocampus—a canonical substrate of SL—may serve computationally distinct roles in SL. Previous work has positioned the CA1 subfield, dominant in anterior hippocampus, as a kind of “mismatch detector” during SL (42, 53, 54, 59, 60), which is fully consistent with the hippocampal prediction errors we observed. Our model comparison analysis showed that these hippocampal signals were best fit as a Bayesian surprise signal derived from an internal memory of the distribution of observed transition events. In contrast, cerebellar error signals were numerically better fit by the delta-rule model. In general, delta-rule prediction errors describe how much a prediction deviates from a learned expectation value whereas our Bayesian model implies that the hippocampus is learning the structure of the environment and responding more when there are outcomes that lead to larger changes in that stored structure. Speculatively, these findings point to a division of labor between these regions during SL, where the hippocampus supports memory-based inference over a learned transition structure while the cerebellum supports incremental trial-by-trial error correction of learned sensory-sensory associations.

This dissociation suggests that the prediction errors we detected in the cerebellum may not be redundant copies of hippocampal prediction errors, but rather the result of a separate process. That said, both areas have been shown to communicate: Our results also show relatively left-lateralized Crus I/II activity in the cerebellum and right-lateralized activity in the hippocampus, consistent with recent functional connectivity studies showing that those two specific regions are significantly functionally correlated (61). More broadly, work in model organisms points to multisynaptic pathways linking the hippocampus and cerebellum (62). If and how cerebello-hippocampal interactions influence SL is thus an important avenue for further research.

Our results are consistent with the general idea that the cerebellum supports implicit forms of learning, both in motor and cognitive domains. Examples of classic cerebellar-dependent motor learning processes, like sensorimotor adaptation and delay eyeblink conditioning (63, 64), are predominantly implicit. Similarly, RL processes, which we recently observed in cerebellar activity (26), can also proceed implicitly (65). In the present study, subjects were not explicitly aware of the statistical regularities in our task. This pattern of results—significant neural correlates of learning but a lack of explicit knowledge—is common in SL (21, 50), supporting the broader idea that cerebellar learning largely operates implicitly.

Although this is the first focused study on cerebellar contributions to unsupervised SL, to our knowledge, prior work can be viewed as consistent with our findings. For example, a recent study showed robust learning and representation of rich temporal statistics in the rodent cerebellum during eyeblink conditioning (66) and other work has identified prediction error-like cerebellar signaling in monkeys when an expected sensory cue is omitted (though such tasks typically required that the monkeys generate motor responses to the unexpected events) (67–69). Moreover, the cerebellum has been implicated in language processing (70) and shows error-like responses to semantically surprising sentences (71), an effect presumably tied to learned statistical regularities. Together, these findings and the results presented here suggest that the cerebellum may track the statistics of the sensory environment across a wide range of contexts, even those that do not require movement control.

Overall, our findings highlight prediction error-based learning as a generalized computation of the cerebellum across both motor and nonmotor contexts alike. Identifying shared cerebellar learning computations across domains could offer clues concerning how cerebellar damage can produce diverse symptoms, from motor dysmetria to wide-ranging social and cognitive deficits. This remains an exciting direction for future investigations.

## Methods

### Participants

Thirty-one individuals participated in the study. We excluded runs with excessive head motion (>2.5mm). We also excluded participants who were not attentive or produced fewer than two usable functional runs. Eleven participants were excluded for excessive motion. Three participants never responded to the cover task so we verified their attention with eye-tracking. Two participants wore glasses that interfered with eye-tracking and were excluded from eye-tracking analyses only. Our final sample included 20 participants (mean age = 23.0 [18-30]; N female = 16). The task protocol was approved by the Yale Institutional Review Board and participants provided informed consent.

### Statistical learning task

The experimental session was two hours long. Participants completed an RL task (26), followed by a visual SL task (**Figure 1a**). Following the functional runs, a T1-weighted anatomical image was collected. The session ended with an explicit test phase for the SL task and a debriefing questionnaire (see Supplemental Information).

Participants completed three runs of the SL task in the scanner. They passively viewed a sequence of abstract fractal stimuli (4 unique fractals per run). Each run consisted of 161 trials (i.e., 160 fractal-to-fractal transitions), with about 40 presentations of each fractal. We manipulated the transitional probability (TP) between stimuli, such that each stimulus was associated with a common transition (0.75 probability) and a rare transition (0.25) (**Figure 1b**). Participants viewed the stimulus shown on the screen for 1s, followed by an ISI. During each run, two stimuli were assigned to short (0.8s) and long (3s) ISIs, respectively.

Participants performed a cover task during the sequences. They were instructed to press a button when a fractal was rotated from its original position (165°). These “probe trials” occurred on 15% of trials in random positions in the sequences unrelated to the transition structure. No feedback was provided.

After the anatomical scan, participants completed a post-scan test (12 trials total) outside of the scanner (**Figure 1c**). On each trial, participants saw four fractals from a scanning run, with one cue stimulus on the top of the screen and three choice stimuli below. They were instructed to choose the stimulus that was most likely to follow the cue stimulus.

### Behavioral analysis

We computed participants’ average d-prime and RTs on the cover task to evaluate whether they were paying attention throughout each run. We also used t-tests to compare the probability of selecting the “common” (0.75) transitions over the other fractals during the post-scan test to examine explicit learning.

### fMRI preprocessing

MR data were acquired with a Siemens 3T Prisma scanner and 64-channel coil at BrainWorks in the Wu Tsai Institute at Yale University and preprocessed using *fMRIPrep* 23.2.1 ((72, 73); RRID:SCR_016216) based on *Nipype* 1.8.6 ((74, 75); RRID:SCR_002502). Details of scan parameters and preprocessing are included in Supplemental Information.

### fMRI analyses

We modeled BOLD signals using general linear models (GLMs) in FSL (version 6.0.7.9). Specific details of nuisance regressors are included in Supplemental Information. To examine transition type prediction errors (i.e., rare > common contrast), we ran a model with boxcar regressors for: (1) button presses, (2) stimuli preceded by short ISIs, (3) stimuli preceded by long ISIs, and (4) ISIs. We also implemented a variant of this model where we split (2) and (3) into separate regressors for the first and second half of trials in order to examine learning. In the stimulus regressors, we coded rare transition trials as 1 and common transition trials as -1 to identify voxels that responded more to surprising transitions. Main results hold when ISI are not separately modeled.

To identify incremental prediction error signals, we used an SL model to simulate trial-by-trial prediction errors for each subject and run. We used a fixed learning rate of .005, as we did not have explicit behavior to fit learning rates for individual participants. The prediction error timeseries were separated by preceding ISI, z-scored, and entered into the GLMs as parametric regressors time-locked to stimulus appearance. The full model included: (1) stimulus appearance, (2) button presses, and (3) onset of ISIs, (4) a parametric regressor with model-based prediction errors for short-ISI trials, and (5) a parametric regressor with model-based prediction errors for long-ISI trials.

We averaged parameter estimates across runs within each participant and performed group-level analyses (FLAME 1+2) with an uncorrected threshold of *p* < 0.001 for whole-brain and cerebellum. We additionally performed ROI analyses on hippocampal subregions and cerebellar lobular ROIs (see Supplemental Information).

### Simulating prediction error timecourses

We formalized SL using a simple forward state transition model to generate prediction error timecourses:

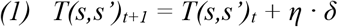

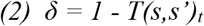

where the transition matrix *T*(*s,s’*) reflects the estimate of the TP from the current stimulus (*s*) to the next (*s’*) on trial *t*, which is updated by the prediction error (δ; difference between the observed transition and the expected transition probability for that stimulus bigram) scaled by learning rate, *η. T* was initialized at 0.25 for all transitions and normalized such that unobserved event probabilities decreased:

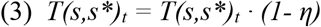

We fixed the learning rate at 0.005 to simulate trialwise SPEs for neural analyses.

### Neural model fitting and comparison

To test whether the cerebellum and hippocampus employ qualitatively different learning algorithms, we fit a delta-rule model and a Bayesian model directly to neural data from Crus I, Crus II and aHPC. BOLD timecourses were extracted from ROIs by running first-level GLMs on each run with only nuisance regressors and stimulus onsets and taking the residualized timeseries. We computed model predictions by generating SPEs at each trial onset and convolving this timeseries with a canonical double-gamma hemodynamic response function. We optimized a single free parameter for each model using *fmincon* in Matlab to minimize the root-mean-square-error between the model-derived and actual timecourses. We repeated parameter optimization 100 times per run to avoid local minima.

The delta-rule model is presented in *Equations 1-3*. The free parameter (*η*) was bounded between [0,1] during fitting.

The Bayesian model stored a matrix, *N*_*t*_*(i,j)*, containing counts of each observed transition from stimulus *I* to the subsequent stimulus *j*. A symmetric Dirichlet prior parameter *a*_*0*_ (pseudo-counts) over all possible transitions was the free parameter of this model (bounded between [0,5] for fitting). Predicted TP for a transition was thus the posterior mean:

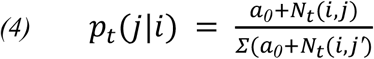

And the prediction error was Bayesian surprise:

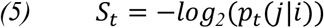

Finally, we computed a **β**-statistic for each model and run using leave-one-run-out cross-validation; we held out one run at a time and used the average of the best-fit parameters (*η* or *a*_*0*_) from the remaining runs to generate the model simulation and regressed that simulation onto the held-out BOLD timecourse.

## Acknowledgements

These data were collected at BrainWorks at the Wu Tsai Institute at Yale University. The authors would like to thank Roeland Hancock and Alex Forrence for technical support at BrainWorks. Additionally, thank you to Tess Levy, Sanghoon Kang, Laurent Caplette, Lily Behm, Diana Wei, Omri Raccah, and Sophie Allen for support with data collection. JET is supported by the NSF GRFP. This project was supported by NIH grant number R01 NS132926 (PI: SDM; Co-PI: NTB).

## Author contributions

S.D.M. procured funding; J.E.T. and S.D.M. designed the research; J.E.T. and Y.O. performed the research; J.E.T. and Y.O. analyzed the data; J.E.T., S.O., N.B.T.-B., and S.D.M. wrote the paper.

