## Supplemental information for "Unsupervised nonmotor learning in the human cerebellum"

### Supplemental Methods

#### *Data acquisition*

Data were collected at BrainWorks in the Wu Tsai Institute at Yale University with a Siemens 3T Prisma scanner and a 64-channel coil. Functional images were collected using multi-band EPI sequences (acceleration factor = 3, interleaved), with TR = 1000 ms, TE = 30 ms, posterior to anterior phase encoding, 48 slices, 2.5 x 2.5 mm in-plane resolution, 3 mm thickness. We collected a high-resolution (1 mm isotropic) T1-weighted (T1w) MPRAGE sequence for anatomical localization and normalization.

#### *Data preprocessing*

Data were preprocessed using *fMRIPrep* 23.2.1 (1, 2); RRID:SCR_016216) based on *Nipype* 1.8.6 (3, 4); RRID:SCR_002502). The T1w image was corrected for intensity non-uniformity (INU) with N4BiasFieldCorrection (5), distributed with ANTs 2.5.0 (6), RRID:SCR_004757), and used as T1w-reference throughout the workflow. The T1w-reference was then skull-stripped with a *Nipype* implementation of the antsBrainExtraction.sh workflow (from ANTs), using OASIS30ANTs as target template. Brain tissue segmentation of cerebrospinal fluid (CSF), white-matter (WM) and gray-matter (GM) was performed on the brain-extracted T1w using fast (FSL (version unknown), RRID:SCR_002823, (7). Brain surfaces were reconstructed using recon-all (FreeSurfer 7.3.2, RRID:SCR_001847, (8), and the brain mask estimated previously was refined with a custom variation of the method to reconcile ANTs-derived and FreeSurfer-derived segmentations of the cortical gray-matter of Mindboggle (RRID:SCR_002438, (9). Volume-based spatial normalization to one standard space (MNI152NLin2009cAsym) was performed through nonlinear registration with antsRegistration (ANTs 2.5.0), using brain-extracted versions of both T1w reference and the T1w template. The following template was were selected for spatial normalization and accessed with *TemplateFlow* (23.1.0, (10): *ICBM 152 Nonlinear Asymmetrical template version 2009c* [(11), RRID:SCR_008796; TemplateFlow ID: MNI152NLin2009cAsym].

The following preprocessing steps were performed for each of the three BOLD runs. First, a reference volume was generated, using a custom methodology of *fMRIPrep*, for use in head-motion correction. Head-motion parameters with respect to the BOLD reference (transformation matrices, and six corresponding rotation and translation parameters) were estimated before any spatiotemporal filtering using mcflirt (FSL; (12). The BOLD reference was then co-registered to the T1w reference using bbregister (FreeSurfer) which implements boundary-based registration (13). Co-registration was configured with 12 degrees of freedom to account for distortions remaining in the BOLD reference. Several confounding time-series were calculated based on the *preprocessed BOLD*: framewise displacement (FD), DVARS and three region-wise global signals. FD was computed using two formulations following Power (absolute sum of relative motions; (14), and Jenkinson (relative root mean square displacement between affines, (12). FD and DVARS are calculated for each functional run, both using their implementations in *Nipype* (following the definitions by (14)). The three global signals are extracted within the CSF, the WM, and the whole-brain masks. Additionally, a set of physiological regressors was extracted to allow for component-based noise correction (*CompCor*, (15)). Principal components were estimated after high-pass filtering the *preprocessed BOLD* time-series (using a discrete cosine filter with 128s cut-off) for the two *CompCor* variants: temporal (tCompCor) and anatomical (aCompCor). tCompCor components are then calculated from the top 2% variable voxels within the brain mask. For aCompCor, three probabilistic masks (CSF, WM and combined CSF+WM) were generated in anatomical space. The implementation differs from that of Behzadi et al. in that instead of eroding the masks by 2 pixels on BOLD space, a mask of pixels that likely contain a volume fraction of GM is subtracted from the aCompCor masks. This mask was obtained by dilating a GM mask extracted from the FreeSurfer's *aseg* segmentation, ensuring that components were not extracted from voxels containing a minimal fraction of GM. Finally, these masks were resampled into BOLD space and binarized by thresholding at 0.99 (as in the original implementation). Components were also calculated separately within the WM and CSF masks. For each CompCor decomposition, the *k* components with the largest singular values were retained, such that the retained components' time series were sufficient to explain 50 percent of variance across the nuisance mask (CSF, WM, combined, or temporal). The remaining components were dropped from consideration. The head-motion estimates calculated in the correction step were also placed within the corresponding confounds file. The confound time series derived from head motion estimates and global signals were expanded with the inclusion of temporal derivatives and quadratic terms for each (16). Frames that exceeded a threshold of 0.5 mm FD or 1.5 standardized DVARS were annotated as motion outliers. Additional nuisance timeseries are calculated by means of principal components analysis of the signal found within a thin band (*crown*) of voxels around the edge of the brain, as proposed by (17). All resamplings can be performed with *a single interpolation step* by composing all the pertinent transformations (i.e. head-motion transform matrices, susceptibility distortion correction when available, and co-registrations to anatomical and output spaces). Gridded (volumetric) resamplings were performed using nitransforms, configured with cubic B-spline interpolation. (Copyright Waiver: The above boilerplate text was automatically generated by fMRIPrep with the express intention that users should copy and paste this text into their manuscripts *unchanged*. It is released under the CC0 license.)

Four participants in the usable sample had one run excluded for excessive motion. The final BOLD data consisted of 56 functional runs across 20 participants. We performed analyses on the whole-brain and separately on masked cerebellar data (as in (18)). The cerebellum was masked on the participant's functional data aligned to standard MNI space and spatially smoothed with a 5-mm kernel. Whole-brain BOLD data were smoothed with a 5-mm kernel before analysis as well.

*Additional GLM analysis methods*

In all GLMs, we extracted motion and noise regressors from the fMRIprep pipeline in addition to the task regressors of interest, including six rigid body regressors, a DVARS regressor to account for overall motion, the first 10 components of the aCompCor regressors to account for noise from white matter or cerebrospinal fluid, and single-TR regressors to scrub high motion TRs (> 0.5 mm). We used the SUIT toolbox (19–21) and nilearn (version 0.8.1) to visualize cerebellar and whole-brain results.

*Additional ROI analysis methods*

Cerebellar ROI masks (Crus I & II) were defined with the Probabilistic Cerebellar Atlas (20). We used the automated segmentation of hippocampus subfields software package (ASHS, Yushkevich et al., 2015) with the ASHS-PMC-T1 atlas (23) to obtain participant-specific ROIs for anterior and posterior Hpc and other MTL subregions (entorhinal cortex (EC); Brodmann areas 35 and 36; parahippocampal cortex (PHC)). We manually inspected each participant to ensure quality. The parahippocampal gyrus ROI (Phg) was defined by combining bilateral EC, BR 35 & 36, and PHC. We transformed MTL ROIs to standard MNI space for analysis.

We extracted averaged **β**-parameters from second-level GLMs within ROIs and conducted nonparametric bootstrap hypothesis tests to test significance. For each ROI, we generated a bootstrap distribution by resampling participants' values with replacement across 1,000 iterations and computing the mean of each resample. Two-tailed p-values were calculated as twice the proportion of bootstrap iterations falling on the side of zero opposite to the mean of the bootstrap distribution. Ninety-five percent confidence intervals were derived from the 2.5th and 97.5th percentiles of the bootstrap distribution.

We conducted ROI analyses to examine responses to transition type and model-inferred prediction errors, as well as to test the temporal specificity of these effects. For transition type, we examined the rare > common contrast (i.e., prediction-error activity) across all trials and separately for short- and long-delay trials. We also examined this effect emerged over time by dividing trials into the first and second half of runs. To examine delta-rule prediction errors, we tested the effect of the parametric regressor across all trials and separately for short- and long-delay trials.

*Physiological analyses*

We used Siemens peripheral devices to collect Pulse and respiration data (same procedure as in (18)). Pulse data were collected by placing a pulse-oximeter on the left index finger. Respiration data were measured with a respiration belt placed around participants' abdomen. Physiological signals were sampled every 2.5ms. We lost a portion of the data due to signal drop-out during the tasks or data storage issues. Of the total usable functional BOLD data (56 runs across 20 participants), the final dataset included 44.6% of the pulse data (25 runs across 11 participants) and 55.4% of the respiration data (31 runs across 13 participants).

For the physiological analysis, we first extracted trial-wise pulse and respiration traces for each run. Then we located the local maxima and minima in the raw physiological signals using the peakdet function in MATLAB (24) and converted the time indices to seconds. We counted the number of maxima and minima that occurred within the interval between stimulus onset of the current trial and the onset of the subsequent trial. We averaged the count of the maxima and minima during this interval (in minutes), since a portion of physiological cycles may not align perfectly with the trial boundaries. Trial-wise pulse and respiration rates were then calculated by dividing the averaged count by the duration of the interval. We averaged these rates separately for rare and common transition trials for each participant and compared them using paired-sample t-tests.

*Post-scan debriefing*

Participants completed a debriefing questionnaire after the post-scan test phase. In the survey, participants were asked to describe any patterns they noticed in the fractal images during the statistical learning task. Most participants (N = 18) focused on describing visual features of the stimuli, such as their shapes, colors, internal lines, or strategies used to judge image orientation. Only two participants specifically reported noticing that the fractals appeared in a particular sequence or order. Of these two participants, one showed a higher-than-chance tendency to select the common transition, whereas the other did not exhibit clear evidence of explicit sequence learning.

### Supplemental Figure

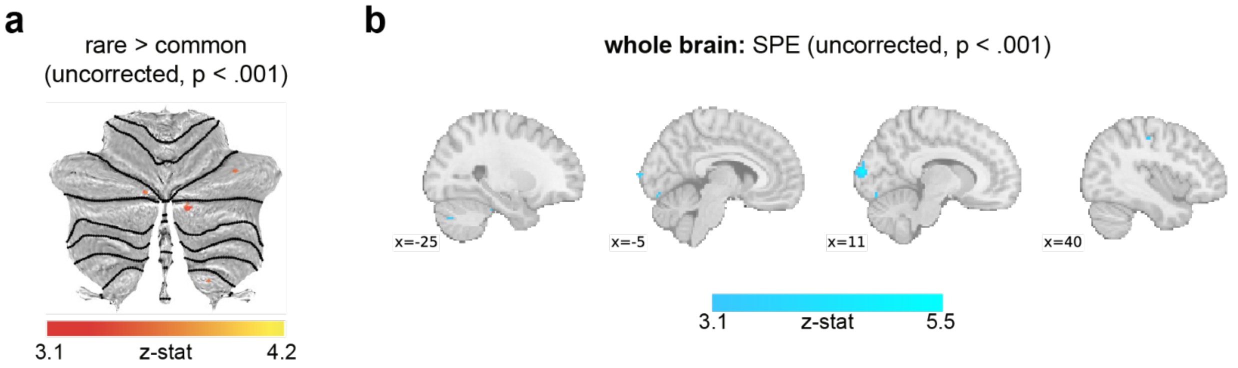


**Supplemental Figure 1. a)** Rare > common effect in cerebellum (uncorrected, $p < .001$). **b)** Whole brain SPE effect (uncorrected, $p < .001$).
